# Oral administration of S-217622, a SARS-CoV-2 main protease inhibitor, decreases viral load and accelerates recovery from clinical aspects of COVID-19

**DOI:** 10.1101/2022.02.14.480338

**Authors:** Michihito Sasaki, Koshiro Tabata, Mai Kishimoto, Yukari Itakura, Hiroko Kobayashi, Takuma Ariizumi, Kentaro Uemura, Shinsuke Toba, Shinji Kusakabe, Yuki Maruyama, Shun Iida, Noriko Nakajima, Tadaki Suzuki, Shinpei Yoshida, Haruaki Nobori, Takao Sanaki, Teruhisa Kato, Takao Shishido, William W. Hall, Yasuko Orba, Akihiko Sato, Hirofumi Sawa

## Abstract

In parallel with vaccination, oral antiviral agents are highly anticipated to act as countermeasures for the treatment of the coronavirus disease 2019 (COVID-19) pandemic caused by severe acute respiratory syndrome coronavirus-2 (SARS-CoV-2). Oral antiviral medication demands not only high antiviral activity but also target specificity, favorable oral bioavailability, and high metabolic stability. Although a large number of compounds have been identified as potential inhibitors of SARS-CoV-2 infection *in vitro*, few have proven to be effective *in vivo*. Here, we show that oral administration of S-217622, a novel inhibitor of SARS-CoV-2 main protease (M^pro^, also known as 3C-like protease), decreases viral load and ameliorates the disease severity in SARS-CoV-2-infected hamsters. S-217622 inhibited viral proliferation at low nanomolar to sub-micromolar concentrations in cells. Oral administration of S-217622 demonstrated eminent pharmacokinetic properties and accelerated recovery from acute SARS-CoV-2 infection in hamster recipients. Moreover, S-217622 exerted antiviral activity against SARS-CoV-2 variants of concern (VOCs), including the highly pathogenic Delta variant and the recently emerged Omicron variant. Overall, our study provides evidence that S-217622, an antiviral agent that is under evaluation in a phase II/III clinical trial, possesses remarkable antiviral potency and efficacy against SARS-CoV-2 and is a prospective oral therapeutic option for COVID-19.

## Main

During the pandemic of the last two years, COVID-19 has caused an increasing number of cases and remains a serious public health concern. Vaccines and antiviral agents for COVID-19 have been developed and some have been approved for clinical use^1^. Oral antiviral medication is highly anticipated to shift the momentum of the pandemic because patients can administer it by themselves and thus benefit from easy access to treatment^2^. To date, two oral bioavailable antivirals have been validated in clinical trials and have been used for the treatment of COVID-19: molnupiravir and nirmatrelvir. Molnupiravir (MPV, also known as EIDD-2801) is a ribonucleoside prodrug of N-hydroxycytidine (NHC) and targets the viral RNA polymerase of SARS-CoV-2. Nirmatrelvir (also known as PF-07321332) is a selective inhibitor of SARS-CoV-2 main protease (M^pro^). The *in vivo* efficacy of both antivirals have been experimentally proven in animal models^3–5^.

SARS-CoV-2 possesses two viral proteases: M^pro^, encoded by the *nsp5* gene, and papain-like protease (PL^pro^), encoded by the *nsp3* gene, which cleave nascent viral polyproteins for maturation in host cells^1^. Because these proteases play essential roles in the intracellular amplification stage of SARS-CoV-2 and lack human homologues, they are ideal targets for specific antivirals^6–8^. S-217622, a novel small-molecule inhibitor for SARS-CoV-2 M^pro^, has been identified through large-scale screening and structure-based optimization^9^ (Fig. 1a). S-217622 showed favorable bioavailability and tolerability in healthy adults in a phase I trial, and has currently been evaluated in a phase II/III trial^10^. Here we report on the antiviral activity of S-217622 against SARS-CoV-2 variants of concern (VOCs) in cell culture and hamsters.

**Fig 1.**
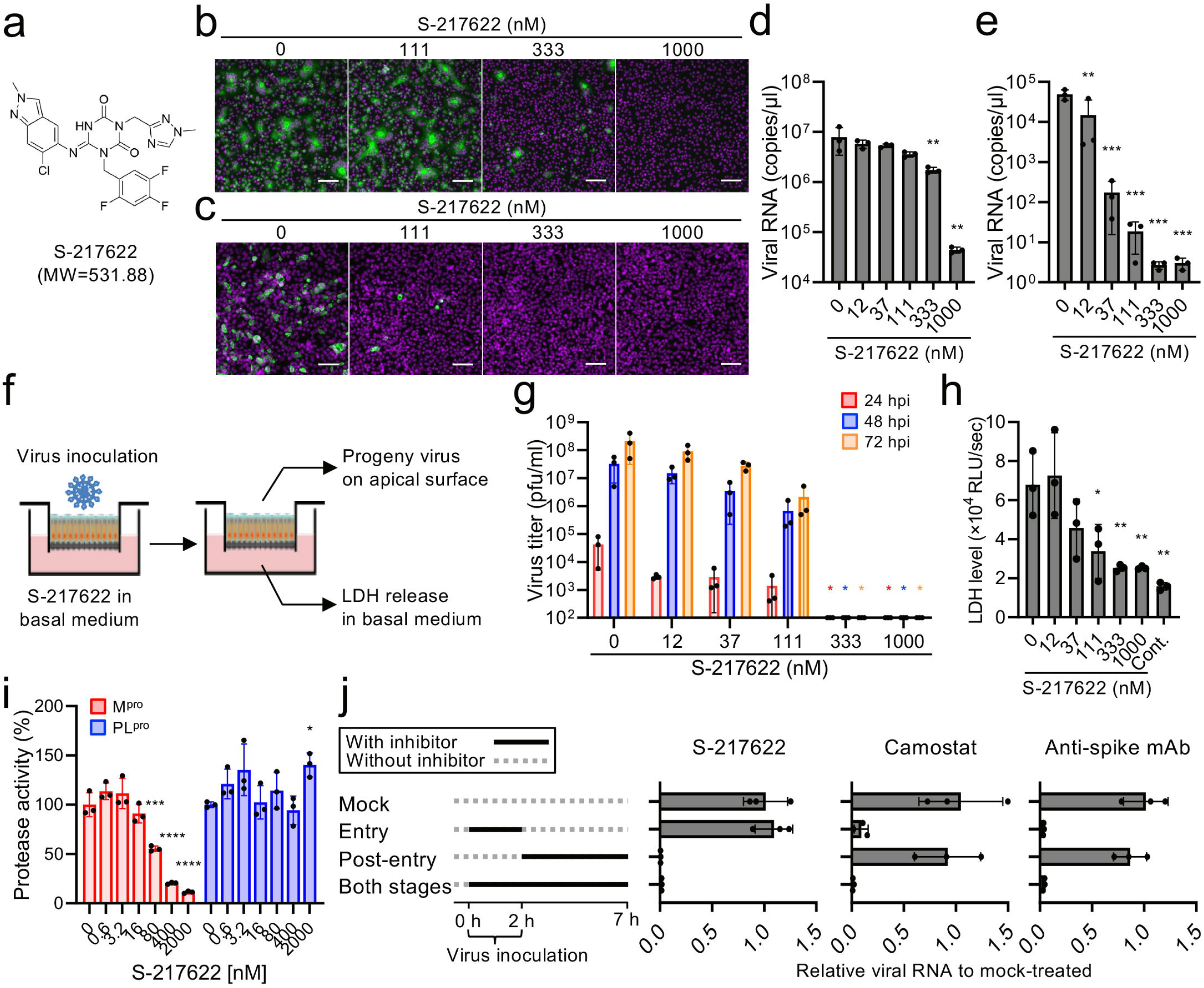
S-217622 inhibits SARS-CoV-2 infection at the post-entry stage. **a**, Chemical structure formula of S-217622. **b**, **c**, Immunofluorescence staining of SARS-CoV-2-infected cells. Vero-TMPRSS2 (**b**) and Calu-3 (**c**) cells were inoculated with a SARS-CoV-2 Delta variant and then cultured in the presence of S-217622 for 24 h and 72 h, respectively. Cells were stained with anti-SARS-CoV-2 nucleocapsid antibody (green) and Hoechst 33342 (magenta). Scale bars, 100 μm. **d**, Viral RNA levels in the culture supernatant of Vero-TMPRSS2 at 24 hours post-infection (hpi) with a SARS-CoV-2 Delta variant. **e**, Viral RNA levels in the culture supernatant of Calu-3 at 72 hpi with SARS-CoV-2 Delta variant. **f**, Schematic representations of the SARS-CoV-2 infection experiment in a human airway tissue model. Viral growth was monitored by titration of progeny virus in the mucus layer on apical surface. The tissue damage by viral infection was estimated by measurement of lactate dehydrogenase (LDH) released from cells into the basal culture medium. **g**, Growth of SARS-CoV-2 Delta variant in a human airway tissue model. Basal culture medium was supplemented with the indicated concentration of S-217622. **h**, The levels of LDH in the basal culture medium was measured by LDH-Glo luminescent assay. **i**, Luciferase-based biosensor assay for SARS-CoV-2 protease activity. 293T cells were co-transfected with expression plasmids of SARS-CoV-2 M^pro^ and a biosensor for SARS-CoV-2 M^pro^ and maintained in the presence of S-217622. The luminescence signal from the biosensor was measured as the protease activity at 24 h post transfection. 293T cells expressing SARS-CoV-2 PL^pro^ and another biosensor for PL^pro^ were used as a control for the target specificity of S-217622. **j**, Time of the addition assay of S-217622. Calu-3 cells were treated with S-217622 during inoculation (entry stage), after inoculation (post-entry stage), or both entry and post-entry stages of SARS-CoV-2. The inhibitory effect of each treatment was evaluated by measurement of intracellular viral RNA level at 7 hpi. Camostat and anti-SARS-CoV-2 spike antibody were used as entry inhibitors for assay controls. The values shown are mean ± standard deviation (SD) of triplicate samples. *p<0.05, **p<0.01, ***p<0.001, ****p<0.0001 by one-way ANOVA with Dunnet’s test (d, e, h, i) or Kruskal-Wallis test with Dunn’s test (g).

### *In vitro* antiviral activity of S-217622 against SARS-CoV-2

To investigate the antiviral activity of S-217622 in cells, we performed cell-based infection assays using the SARS-CoV-2 Delta variant (lineage B.1.617.2). The infection with SARS-CoV-2 was inhibited in Vero-TMPRSS2 and Calu-3 cells by S-217622 in a dose-dependent manner (Figs. 1b, c). S-217622 treatment also dose-dependently reduced the progeny viral load in the culture supernatants from both Vero-TMPRSS2 and Calu-3 cells (Figs. 1d, e).

Primary human bronchial epithelial cells (HBE cells) can be differentiated under air-liquid interphase culture conditions and used as an *ex vivo*model for SARS-CoV-2 infection^11^. HBE cells were exposed to SARS-CoV-2 and treated with S-217622 though the basal medium (Fig. 1f). The progeny virus titers were decreased at low concentrations (12, 37, and 111 nM) and were under the detection limit of the plaque assay (<100 plaque forming unit/ml) at 333 nM or more of S-217622 (Fig. 1g). Lactate dehydrogenase (LDH) release into basal culture medium is an indicator of cytotoxicity by virus infection and was decreased in the presence of S-217622 (Fig. 1h). The antiviral potency of S-217622 was similar to or higher than those of the antivirals approved for clinical use (nirmatrelvir, molnupiravir, and remdesivir; Extended Data Table 1). Since nirmatrelvir and remdesivir are major substrates for the plasma membrane multidrug transporter P-glycoprotein (P-gp), these antivirals require a P-gp inhibitor such as CP-100356 to inhibit the efflux of the antivirals from Vero cells, which express high levels of P-gp^12^. These data indicate that S-217622 inhibits SARS-CoV-2 infection at nanomolar to submicromolar concentrations in different cells.

To date, seven variants, namely, Alpha, Beta, Gamma, Delta, Omicron, Lambda, and Mu have emerged and have been assigned as VOC or variants of interest (VOI) by WHO. These variants possess amino acid changes in the spike protein, altering the infectivity, transmissibility, pathogenicity, or susceptibility to antibody neutralization^13–16^. Unlike the S gene, which encodes the viral spike protein, the *nsp5* gene, which encodes M^pro^, a target for S-217622, is well conserved among these VOCs and VOIs (Extended Data Fig. 1). Indeed, S-217622 exhibited antiviral activity against all VOCs, while neutralizing antibodies had different reactivities to some VOCs due to mutations in the spike protein (Extended Data Figs. 2a, b)^15, 17^. The remarkable antiviral potency of S-217622 was also reproduced in Vero-TMPRSS2 cells infected with the SARS-CoV-2 Omicron variant (lineage BA.1) (Extended Data Figs. 3a, b).

### S-217622 disrupts the post-entry stage of SARS-CoV-2 infection

To validate the antiviral mechanism of S-217622, we employed a luciferase-based biosensor assay for measurement of SARS-CoV-2 M^pro^ activity^18^. GloSensor-AVLQS contains a M^pro^-cleavage motif and acts as an indicator of the protease activity of M^pro^. GloSensor-RLKGG is a substrate of another SARS-CoV-2 protease, PL^pro^, and was used as a control. S-217622 inhibited the cleavage/activation of GloSensor-AVLQS by M^pro^ but not the cleavage/activation of GloSensor-RLKGG by PL^pro^, indicating the target specificity of S-217622 (Fig. 1i). The time of addition assay showed that S-217622 has an effect on the post-entry process but not the cellular entry process of infection (Fig. 1j). These data suggest that S-217622 blocks SARS-CoV-2 infection through the inhibitory effect of viral M^pro^ activity at a post-entry stage.

### Prophylactic administration of S-217622 in hamsters

We investigated the *in vivo* antiviral activity of S-217622 using Syrian hamsters in an animal model for COVID-19^19, 20^. The pharmacokinetic parameters in plasma revealed the excellent bioavailability of S-217622 (Extended Data Table 2). After oral administration of S-217622, its concentration in plasma reached the maximum at 0.5 to 2.67 h after doses of 10, 30, and 100 mg/kg in hamsters, and then declined with the t_1/2,z_ values of 3.43 to 4.46 h (Fig. 2a and Extended Data Table 2). The C_max_ and AUC were increased more than the dose ratio at 10 to 100 mg/kg. Based on the dose-dependency in hamsters, we set two different dose levels (30 mg/kg and 200 mg/kg) for our *in vivo* experiments.

**Fig 2.**
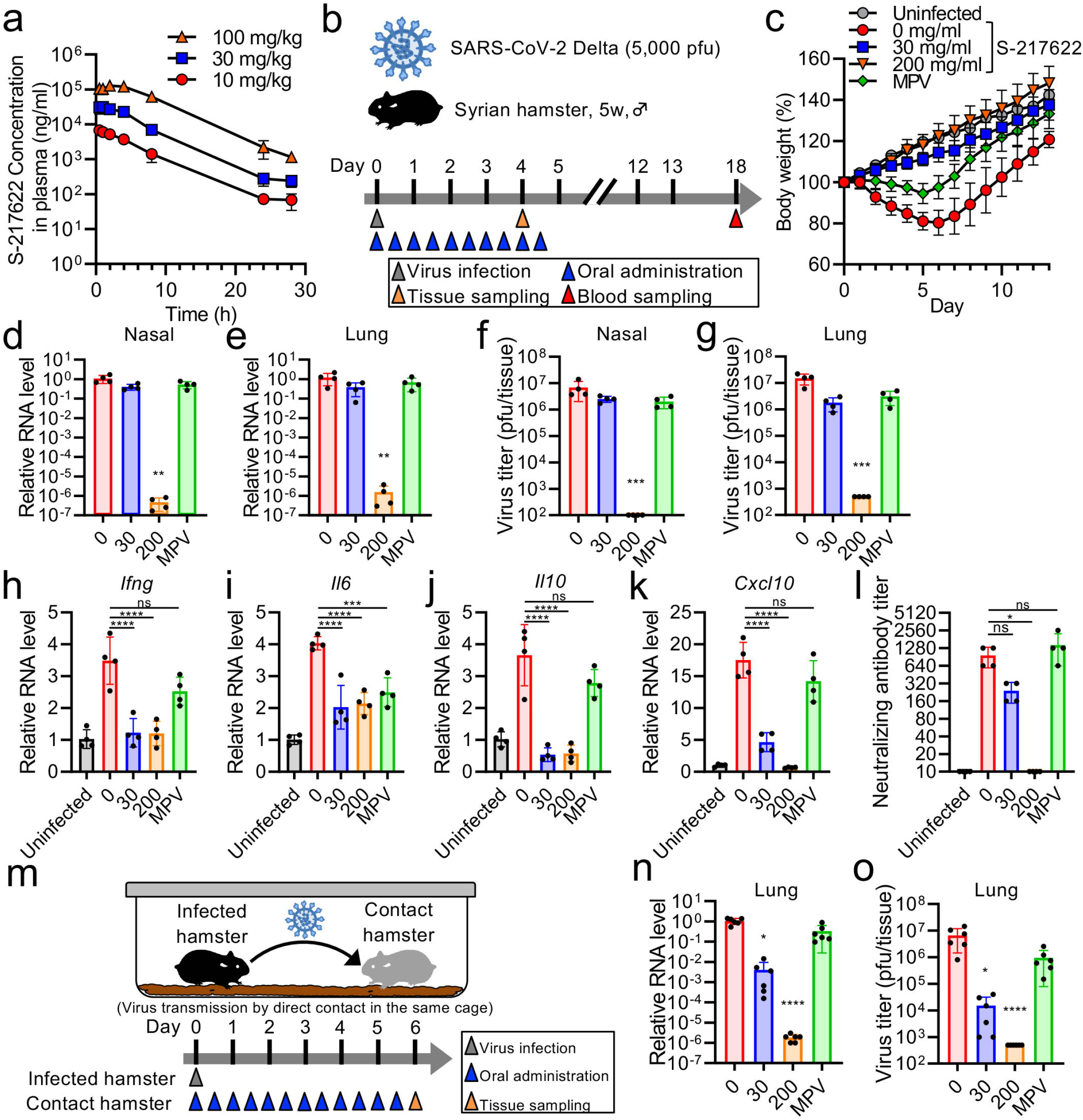
Prophylactic treatment of S-217622 prevents viral growth and the onset of COVID-19 in hamsters inoculated with SARS-CoV-2. **a**, Plasma concentration profile of S-217622 after a single oral administration in hamsters. Plasma samples were harvested at the indicated time points and analyzed by LC/MS/MS. **b**, Schematic of the experimental design for prophylactic treatment in a hamster model. Hamsters were intranasally inoculated with 5,000 plaque-forming units (pfu) of SARS-CoV-2 Delta variant. For prophylactic treatment, the hamsters were treated with oral administration of S-217622 or vehicle (0 mg/ml) twice daily (b.i.d.) from the time of infection to 4 dpi. Molnupiravir (MPV) was used as a comparator drug. A group of hamsters were sacrificed at 4 dpi for tissue collection. **c**, Body weight changes in uninfected hamsters (n = 3) and SARS-CoV-2-infected hamsters with the treatments of S-217622 (30 mg/ml and 200 mg/ml), vehicle (0 mg/ml), or MPV (200 mg/kg) (n = 4 for each group). **d**, **e**, Viral RNA levels in nasal turbinates (**d**) and lungs (**e**) from hamsters at 4 dpi with SARS-CoV-2. Each group of hamsters was treated with vehicle (red), 30 mg/kg (blue), and 200 mg/kg (orange) of S-217622 or MPV (green). Relative viral RNA levels in lungs as compared with lungs from vehicle-treated hamsters were examined. Data were normalized to β-actin. **f**, **g**, Virus titers in nasal turbinates (**f**) and lungs (**g**) from hamsters at 4 days post-infection (dpi) with SARS-CoV-2 were determined by plaque assay. Each group of hamsters was treated with vehicle (red), 30 mg/kg (blue) and 200 mg/kg (orange) of S-217622, or MPV (green). **h**-**k**, Cytokine gene expression profiles in lungs from hamsters at 4 dpi with SARS-CoV-2. Relative gene expression levels of indicated cytokines in the lungs as compared with lungs from uninfected hamsters were examined using qRT-PCR. Data were normalized to β-actin. l, Neutralizing antibody titers in hamster serum at 18 dpi. **m**, Schematic of the experimental design for virus transmission from infected hamster to naïve hamster. One hamster per cage was inoculated with 5,000 pfu of SARS-CoV-2 Delta variant (infected hamster). Three naïve hamsters (contact hamsters) were co-housed with the infected hamster in the same cage. Only the contact hamsters were prophylactically treated with S-217622 or MPV (b.i.d.) **n**, **o**, Viral RNA levels (**n**) and virus titers (**o**) in lungs from the contact hamsters after 6 days of co-housing with the infected hamster. Each group of the contact hamsters was treated with vehicle (red), 30 mg/kg (blue) and 200 mg/kg (orange) of S-217622, or MPV (green). For the comparison of viral RNA levels, relative viral RNA levels in lungs compared to lungs from vehicle-treated hamsters were examined. Data were normalized to -actin. The values shown are mean ± SD. ns = not β significant, *p<0.05, **p<0.01, ***p<0.001, ****p<0.0001 by Kruskal-Wallis test with Dunn’s test (d-g, l, n, o) or one-way ANOVA with Tukey’s test (h-k).

Hamsters were inoculated with SARS-CoV-2 Delta, a highly pathogenic variant^13^, followed by oral administration of antivirals twice a day (b.i.d.) from the time of inoculation (Fig. 2b). Vehicle control hamsters (0 mg/kg) were orally administered with the vehicle only. We employed molnupiravir (MPV, dose = 200 mg/kg) as a comparator drug, which has been reported to reduce the viral load of SARS-CoV-2 in hamsters^5^. Control hamsters lost more than 15% of body weight at 4–7 days post-infection (dpi), while hamsters treated with MPV showed only 5% body weight loss until 6 dpi (Fig. 2c). In contrast, hamsters treated with S-217622 continuously gained the weight through the experimental period. Viral RNA loads and titers in nasal turbinates and lungs of hamsters treated with antivirals were lower than that of vehicle group (Figs. 2d-g). Notably, treatment with S-217622 (200 mg/kg) resulted in a more than 10^5^-fold decrease in viral RNA load and the titers were under the detection limit in hamsters at 4 dpi (Figs. 2f, g).

The antiviral activity of S-217622 was also observed in hamsters infected with other VOCs: Alpha (lineage B.1.1.7), Gamma (lineage P.1), and Omicron (Extended Data Figs. 4a-f). SARS-CoV-2 infection caused severe pneumonia and induced host inflammatory responses in hamsters, while treatment with S-217622 decreased the expression levels of inflammatory cytokines (Figs. 2h-k). Hosts develop specific antibody from 10 to 14 dpi with SARS-CoV-2^21^. Notably, no seroconversion was observed in hamsters treated with S-217622 (200 mg/kg) (Fig. 2l). These results suggest that S-217622 has antiviral activity *in vivo* and that prophylactic administration protects hamsters from SARS-CoV-2 infection and the onset of COVID-19.

We examined whether post-exposure prophylactic administration of S-217622 prevented viral transmission to susceptible hamsters (contact hamster) by co-housing susceptible hamsters with SARS-CoV-2-infected hamsters in the same cage (Fig. 2m). Susceptible hamsters continued to be exposed to virus shed from SARS-CoV-2-infected hamsters. After 6 days of co-housing with SARS-CoV-2-infected hamsters, high viral load was detected in the lungs of the contact hamsters treated with vehicle or MPV (Figs. 2n, o). However, prophylactic administration of S-217622 (200 mg/kg) resulted in a more than 10^5^-fold reduction in viral RNA load and under the detection limit of virus titer in the lung of hamsters at 4 dpi. The results suggested that prophylactic administration of S-217622 can inhibit viral spread among co-housed hamsters, highlighting the high level of *in vivo* antiviral efficacy of S-217622.

### Therapeutic administration of S-217622 in hamsters

SARS-CoV-2 replicates rapidly in hamsters intranasally inoculated with SARS-CoV-2, and the viral load in the upper respiratory tract reaches its peak at 1–2 dpi^13, 22^. To evaluate the therapeutic efficacy of S-217622, we treated hamsters with S-217622 from 1 dpi (delayed treatment protocol) (Fig. 3a). The vehicle and MPV groups showed similar body weight curves and lost more than 10% of body weight at 5–6 dpi (Fig. 3b). In contrast, the peak of body weight loss in hamsters receiving S-217622 was 7% (30 mg/kg) and 5% (200 mg/kg) and body weight recovery was accelerated by the administration of S-217622.

**Fig 3.**
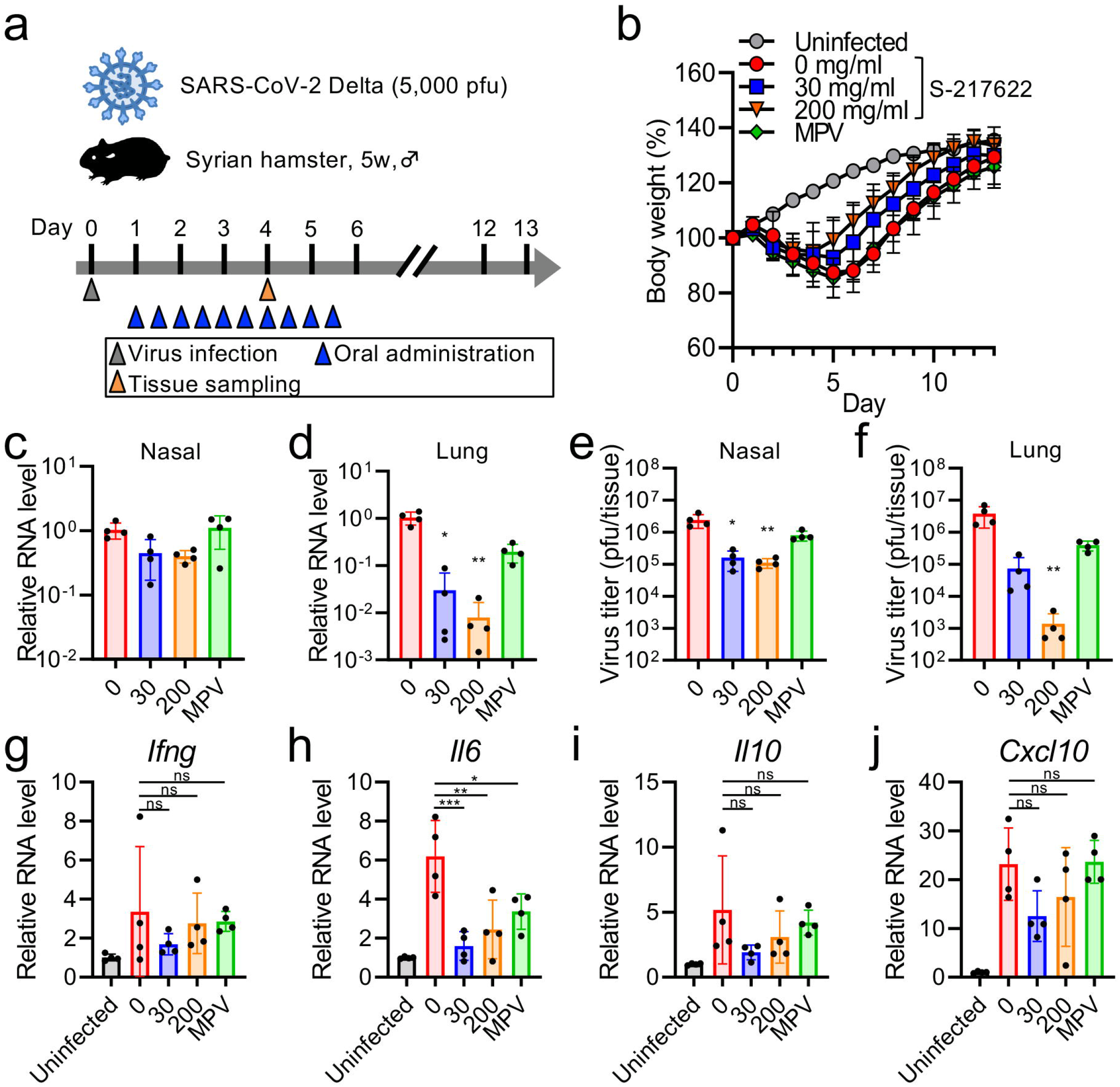
Therapeutic treatment with S-217622 decreased the viral load of SARS-CoV-2 and disease severity in hamsters. **a**, Schematic representation of the experimental design for therapeutic treatment in a hamster model. Hamsters were intranasally inoculated with 5,000 pfu of a SARS-CoV-2 Delta variant. For therapeutic treatment, the hamsters were treated with oral administration of S-217622 or vehicle twice daily from 1 dpi to 5 dpi. Molnupiravir (MPV) was used as a comparator drug. A group of hamsters were sacrificed at 4 dpi for tissue collection. **b**, Body weight change in uninfected hamsters (n = 3) and SARS-CoV-2-infected hamsters with the treatments of S-217622 (30 mg/ml and 200 mg/ml), vehicle (0 mg/ml), or MPV (n = 4 for each group). **c**, **d**, Viral RNA levels in nasal turbinates (**c**) and lungs (**d**) from hamsters at 4 dpi with SARS-CoV-2. Each group of hamsters was treated with vehicle (red), 30 mg/kg (blue) and 200 mg/kg (orange) of S-217622, or MPV (green) from 1 dpi. Relative viral RNA levels in lungs as compared with levels in lungs from vehicle-treated hamsters were examined. Data were normalized to β-actin. **e**, **f**, Virus titers in nasal turbinates (**e**) and lungs (**f**) from hamsters after 4 dpi with SARS-CoV-2 were determined by plaque assay. Each group of hamsters was treated with vehicle (red), 30 mg/kg (blue) and 200 mg/kg (orange) of S-217622, or MPV (green) from 1 dpi. **g**-**j**, Cytokine gene expression profiles in lungs from hamsters at 4 dpi with SARS-CoV-2. Relative gene expression levels of indicated cytokines in the lungs as compared with lungs from uninfected hamsters were examined using qRT-PCR. Data were normalized to β-actin. The values shown are mean ± SD. ns = not significant, *p<0.05, **p<0.01, ***p<0.001 by Kruskal-Wallis test with Dunn’s test (c-f) or one-way ANOVA with Tukey’s test (g-j).

Administration of S-217622 or MPV had a limited effect on viral loads in nasal turbinates, but these antivirals significantly decreased the viral loads in the lungs at 4 dpi (Figs. 3c-f). Administration of S-217622 (200 mg/kg) achieved a greater than 10^3^-fold decrease of virus titer in the lung of hamsters at 4 dpi (Fig. 3f). Administration of S-217622 decreased the expression levels of inflammatory cytokines in part, and the contribution to the control of the host inflammatory response was limited (Figs. 3g-j). We sequenced the *nsp5* gene and found no homogenous amino acid substitution in progeny viruses in the lungs from S-217622-treated hamsters. These results suggest that administration of S-217622 decreases the viral load in the lungs and facilitates recovery from the infection, even with delayed treatment.

### Histopathological findings in the lungs of SARS-CoV-2-infected hamsters

In the prophylactic experiments (Fig. 2b), histopathological findings of viral pneumonia were prominent among the vehicle group and the MPV-administered hamsters (Fig. 4a). In contrast, S-217622 attenuated the lung pathology in a dose-dependent manner (Fig. 4a), and the histopathological severity score of the S-217622-administered (200 mg/kg) group was significantly lower than the vehicle control group (Fig. 4b). Viral RNA was clearly decreased following both MPV and S-217622 (30 mg/kg) administration (Fig. 4a). It was remarkable that viral RNA was hardly detected by *in situ* hybridization (ISH) in the S-217622 (200 mg/kg)-administered hamsters (Fig. 4a), consistent with the results of qRT-PCR (Fig. 2e).

**Fig 4.**
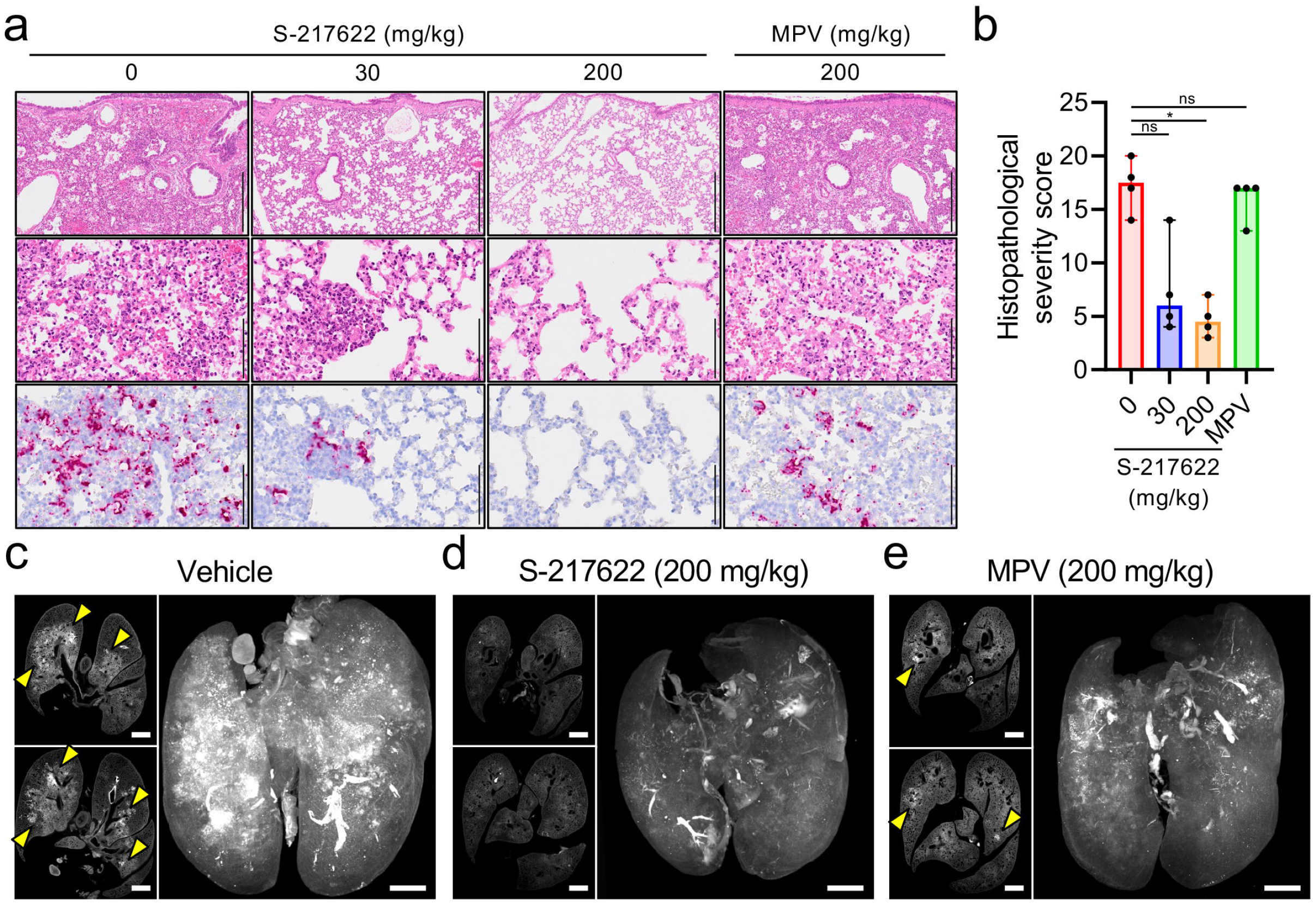
Histopathological findings in the lungs of SARS-CoV-2-infected hamsters that were administered antivirals. The hamsters were infected with the SARS-CoV-2 Delta variant and sacrificed at 4 dpi for histopathological examinations. S-217622 (30 mg/kg or 200 mg/kg), molnupiravir (200 mg/kg), or vehicle control (0 mg/ml) was administered from 0 to 3 dpi (**a**, **b**) or 1 to 3 dpi (**c**-**e**) twice a day following the schedules shown in Fig. 2b and 3a, respectively. **a**, Representative histopathological images for the lung sections obtained from the animals given antivirals indicated above each panel (n = 4). Upper and middle panels, hematoxylin and eosin (H&E) staining. Lower panels, *in situ* hybridization (ISH) targeting the nucleocapsid gene of SARS-CoV-2. Scale bars in upper panels, 500 μm. Scale bars in middle and lower panels, 100 μm. **b**, Histopathological severity score of pneumonia based on the percentage of alveolitis area in a given section. Data are shown as the median score ± 95% confidential interval with each dot representing the score of each animal. (n = 4; ns = not significant, *p<0.05 by Kruskal-Wallis test with Dunn’s test.) **c**-**e**, Cross-sectional imaging of lungs and 3D image reconstruction of whole lungs from hamsters at 4 dpi. The infected hamsters were treated with vehicle (c), S-217622 (d), or MPV (e). The whole lung tissues were stained with anti-SARS-CoV-2 spike antibody and scanned by light sheet microscopy. Arrowheads indicate foci of SARS-CoV-2-positive alveoli. Scale bars, 2 mm.

In the therapeutic experiments, histopathological examination of the lungs of vehicle control hamsters revealed massive infiltration of inflammatory cells, including neutrophils and lymphocytes, into alveoli and the alveolar walls accompanied with alveolar hemorrhage (Extended Data Fig. 5a). These findings were also detected in the drug-administered hamsters (Extended Data Fig. 5a), and neither MPV or S-217622 could attenuate the histopathological severity of viral pneumonia by therapeutic treatment (Extended Data Fig. 5b). Viral RNA was readily detected by ISH across extended areas of the lungs from the vehicle group (Extended Data Fig. 5a). Positive signals for viral RNA were slightly decreased in the MPV-administered animals, while a significant reduction was observed in the S-217622-administered groups (Extended Data Fig. 5a), which showed a discrepancy with the finding that the histopathological severity of viral pneumonia was similar between the two groups.

The viral antigen distributions in the lungs of the therapeutic administered groups were investigated using whole-imaging approaches with light sheet microscopy. Whole lung tissues were fixed, bleached, and stained with anti-SARS-CoV-2 antibody and then exposed to clearing solution for scanning. Slice images showed bronchial and peribronchial distribution of viral antigen in vehicle-treated hamsters (Fig. 4c). Reconstructed 3D images revealed that virus infection spread throughout the lungs of the vehicle-treated controls (Fig. 4c and Supplementary Video 1). Consistent with the results of ISH, the signal distribution of viral antigen was limited in the lungs of hamsters treated with S-217622 (Fig. 4d and Supplementary Video 2). Administration of MPV decreased the viral distribution but the efficacy was relatively limited compared to administration of S-217622 (Fig. 4e and Supplementary Video 3). Taken together, these findings suggested that SARS-CoV-2 replication was significantly inhibited by both prophylactic and therapeutic administration of S-217622, and that the progression of lung pathology due to viral pneumonia was also suppressed by prophylactic administration of S-217622.

## Discussion

In this study, we characterized the antiviral activity of S-217622, a newly identified SARS-CoV-2 M^pro^ inhibitor. Our *in vitro* experiments revealed that S-217622 exhibits remarkable antiviral potency against all VOCs. Current vaccines and monoclonal antibody medications target the viral spike protein, which accumulates amino acid variations among SARS-CoV-2 variants^16, 17^. In contrast, viral M^pro^ is less divergent, leading to the comparable susceptibility of VOCs to S-217622. Although host factors involved in virus proliferation could be targets for antivirals, viral proteins are specific and favorable targets for antiviral development^23^. Overall, SARS-CoV-2 M^pro^ is a prospective target and the inhibitor S-217622 is a broadly reactive antiviral against SARS-CoV-2 variants.

Prophylactic administration of S-217622 prominently decreased viral load in both nasal turbinates and lung tissues. Hamsters receiving high doses of S-217622 developed no detectable neutralizing antibodies at 18 days after inoculation. This result indicates that administration of S-217622 confers sterile protection against SARS-CoV-2 in the recipient animal and suggests the potential of S-217622 as a preventive medication for high-risk individuals who have close contact with patients with COVID-19, and potential treatment of those with active disease. In animal experiments, prophylactic administration at pre-infection or infection stages is a highly sensitive experimental method to detect the efficacy of antivirals^24, 25^. Our experiments showed that S-217622 has an anti-SARS-CoV-2 effect and facilitates recovery from the acute infection stage even in a post-exposure treatment environment, highlighting the high antiviral activity and the therapeutic potential of S-217622.

MPV and nirmatrelvir have been approved as oral antiviral medications for clinical use; however, concerns remain regarding the efficacy and potential risks. The results of a recent clinical trial showed that MPV treatment decreased the risk of hospitalization for COVID-19 by 30%^26, 27^. Our experiments also showed that therapeutic treatment with MPV had a limited effect on viral load and body weight decrease in hamsters. It also has been reported that MPV has mutagenesis for not only viral RNA but also host DNA in cell-based assays^28^. Nirmatrelvir is another M^pr^° inhibitor and is reported to have an excellent *in vivo* efficacy, with an 88% reduction in the risk of hospitalization or death^29^. A previous study, along with our data, shows that nirmatrelvir is highly sensitive to the multidrug transporter, P-gp, and requires a P-gp inhibitor to exhibit activity in Vero cells^12^. For clinical use, nirmatrelvir requires co-administration with a CYP3A4 inhibitor (ritonavir) as a pharmacokinetic booster to slow the metabolism of nirmatrelvir. Because the ritonavir booster also affects the metabolism of other medications, clinicians need to consider the potential other drug-drug interactions during treatments^30^. In contrast, S-217622 showed expected bioavailability and concentration without any pharmacokinetic booster for humans in a phase I trial^10^. Consequently, S-217622 has different biological properties from the preceding medications and is expected to be an alternative candidate for COVID-19 therapy.

We note some limitations of our study. First, therapeutic treatment with S-217622 was insufficient to control pneumonia in SARS-CoV-2-infected hamsters, although they regained lost body weight earlier compared to vehicle controls. We assume that the initial virus proliferation stimulated host immunity and subsequent inflammation^31, 32^, and combination with antiviral and anti-inflammatory medication may lead to better results and outcomes^33^. Second, this study was conducted using hamsters as a COVID-19 model and the efficacy in human patients cannot be inferred. However, S-217622 is currently under evaluation in a phase II/III clinical trial. Third, the development of a resistant viral clone against S-217622 and its virological properties will need to be investigated in future studies.

In summary, our study has demonstrated the remarkable antiviral activity of S-217622 through *in vitro* and *in vivo* experiments. This scientific evidence will be invaluable when considering the application of S-217622 as a medication for COVID-19.

## Supporting information

Supplemental Video S1, related to Fig. 4c

Supplemental Video S2, related to Fig. 4d

Supplemental Video S3, related to Fig. 4e

## Methods

### Ethics statement

The animal experiments with virus infection were performed in accordance with the National University Corporation, Hokkaido University Regulations on Animal Experimentation. The protocol was reviewed and approved by the Institutional Animal Care and Use Committee of Hokkaido University (approval no. 20-0060). The animal experiments for pharmacokinetics analysis were approved by the Director of the institute after reviewing the protocol by the Institutional Animal Care and Use Committee in SHIONOGI & CO., Ltd. (approval no. S21187C-0001).

### Cells

Calu-3 cells (ATCC, HTB-55) were maintained in Eagle’s Minimum Essential Medium (MEM) containing 10% fetal bovine serum (FBS). Vero-TMPRSS2 cells [Vero E6 cells (ATCC, CRL-1586) stably expressing human TMPRSS2]^34^ were maintained in Dulbecco’s Modified Eagle’s Medium (DMEM) containing 10% FBS. 293T (RIKEN BRC, RCB2202) cells were maintained in high glucose DMEM containing 10% FBS.

293T-ACE2-TMPRSS2 cells stably expressing human TMPRSS2 and ACE2 were established by a lentiviral vector transduction system as previously described^35^ and maintained in high glucose DMEM containing 10% FBS. Differentiated human bronchial epithelial cells were obtained as MucilAir-bronchial (Epithelix, EP01) and maintained under an air-liquid interphase culture condition with MucilAir culture medium (Epithelix) according to manufactures’ instruction.

### Viruses

SARS-CoV-2 VOC Alpha (strain QK002, lineage B.1.1.7, GISAID: EPI_ISL_768526), Beta (strain TY8-612, lineage B.1.351, GISAID: EPI_ISL_1123289) , Gamma (strain TY7-501, lineage P.1, GISAID: EPI_ISL_833366) , Delta (strain TY11-927, lineage B.1.617.2, GISAID: EPI_ISL_2158617) , Omicron (strain TY38-873, lineage BA.1, GISAID: EPI_ISL_7418017) were provided from National Institute of Infectious Diseases, Japan. The working viral stocks were prepared by a passage on Vero-TMPRSS2 cells. The titers of the prepared virus stocks were determined as plaque forming unit per ml (pfu/ml) by a plaque assay.

### Compounds

S-217622 (fumaric acid co-crystal form) and Nirmatrelvir were synthesized by Shionogi & Co., Ltd.^9^. MPV and remdesivir were obtained from MedChemExpress. NHC was obtained from Angene. REGN10933 and REGN10987 were obtained from Cell Sciences.

### Plaque assay

Monolayers of Vero-TMPRSS2 were inoculated with serial dilutions of either virus stock or experiment samples for 1h at 37°C. The cells were then overlaid with DMEM containing 2% FBS, 0.5% Bacto Agar (Becton Dickinson) and 25 μ (Wako). At 3 dpi with omicron and 2 dpi with other variants, cells were fixed with 3.7% formaldehyde in PBS and stained with 1% crystal violet.

### Immunofluorescence staining

Vero-TMPRSS2 and Calu-3 cells were inoculated with SARS-CoV-2 at multiplicities of infection (MOIs) of 0.1 and 10, respectively. After 1h incubation, cells were fed with fresh culture medium containing 2% FBS and S-217622. Vero-TMPRSS2 cells at 24 hpi and Calu-3 cells at 72 hpi were fixed with 3.7% formaldehyde in PBS, followed by permeabilization with 0.5% Triton X-100 in PBS for 5 min and staining with anti-SARS-CoV-2 nucleocapsid rabbit monoclonal antibody (GTX635679, GeneTex) in 25% Block Ace (KAC) in PBS for 1 h. Alexa Fluor 488-conjugated anti-rabbit IgG antibody (Invitrogen; Thermo Fisher Scientific) was used as the secondary antibody. Nuclei were stained with Hoechst 33342 (Invitrogen). Fluorescent images were captured using IX73 fluorescence microscope (Olympus).

### qRT-PCR

Vero-TMPRSS2 and Calu-3 cells were inoculated with SARS-CoV-2 at MOIs of 0.01 or 0.1, respectively. After 1 h of incubation, the cells were washed three times with phosphate-buffered saline (PBS) and fed with fresh culture medium containing 2% FBS and S-217622. The culture supernatants were harvested at 24 hpi from Vero-TMPRSS2 cells and 72 hpi from Calu-3 cells. Viral RNA in the supernatant was extracted with High Pure Viral RNA kit (Roche). The viral copy number was estimated by qRT-PCR with Thunderbird Probe One-step qRT-PCR Kit (Toyobo). Probe and primers targeting SARS-CoV-2 *N* gene were previously described as N2 set^36^.

For measurement of host gene expression and viral RNA levels in hamsters, total RNA was extracted from tissue homogenates with a combination of TRIzol LS (Invitrogen) and Direct-zol RNA MiniPrep kit (Zymo Research). For relative quantification of viral RNA and host mRNAs, RNA samples were analyzed by qRT-PCR with Thunderbird Probe One-step qRT-PCR Kit. Target RNA levels were normalized to hamster -actin and β calculated by the ΔΔCt method. Primers and probes for SARS-CoV-2 *N* gene were noted above. Other primers and probes for *Actb*, *Ifng*, *Il6*, *Il10*, *Cxcl10* genes ^37^ were previously described.

### Evaluation of antiviral efficacy S-217622 in human bronchial epithelial cells

The apical area of human bronchial epithelial cells were washed with culture medium and then inoculated with 5,000 pfu of SARS-CoV-2. After 30 min of incubation, the apical area was washed with PBS and the basal medium was replaced with fresh culture medium supplemented with S-217622. At 24, 48, and 72 hpi, 100 μl of culture medium was added at μl the apical area and harvested for virus titration after 20 min incubation. At 72 hpi, the level of LDH in the basal culture medium was quantified by LDH-Glo Cytotoxicity Assay kit (Promega).

### Biosensor assay for viral protease activity

The DNA fragments encoding SARS-CoV-2 NSP3 or NSP4, NSP5, N-terminal NSP6 (NSP4/5/6N) were cloned into pCMV-derivative pCXSN vector to generate pSARS-CoV-2 PL^pro^ and pSARS-CoV-2 M^pro^, respectively. The amino acid sequence AVLQS (for the cleavage by M^pro^) or RLKGG (for the cleavage by PL^pro^) were inserted into pGloSensor-30F vector backbone (CS182101, Promega) to generate firefly luciferase-based biosensor expressing plasmid pGS-AVLQS or pGS-RLKGG, respectively. 293T cells on 96-well plate were co-transfected with a set of pGS-AVLQS and pSARS-CoV-2 M^pro^ or a set of pGS-RLKGG and pSARS-CoV-2 PL^pro^ using TranIT-LT1 (Mirus Bio) in the presence of serial diluted S-217622. At 24 h post transfection, the luminescence signals of biosensors and renilla luciferase (an internal control reporter) from pGS vectors were measured by Dual-Glo Luciferase Assay System (Promega). The ratio (firefly luciferase/renilla luciferase) was calculated to normalize for the influence of transfection efficiency. Non-treated cells were used as a control for 100% biosensor activation.

### Time of addition assay

Calu-3 cells were inoculated with SARS-CoV-2 Delta variant at an MOI of 3 for 2 h in M), camostat (50 μM, Wako) or anti-SARS-CoV-2 spike (1 μg/ml, GTX635792, GeneTex)]. After washing three times with PBS, cell were cultured for 5 h with or without inhibitors. Total RNA was extracted from the cells using Purelink RNA Mini kit (Invitrogen) and subjected to measurement of intracellular viral RNA levels by qRT-PCR. Target RNA levels were normalized to human β-actin (Hs99999903_m1, Applied Biosystems; Thermo Fisher Scientific) and calculated by the ΔΔCt method. Primers and probes for SARS-CoV-2 *N* gene were as noted above.

### Cytopathic effect-based cell viability assays

Antiviral compounds and antibodies were serially diluted 2-fold increments by culture medium containing 2% FBS and plated on 96-well microplates. The diluted compounds in the plates were mixed with SARS-CoV-2 and cell suspension. The diluted antibodies were initially incubated with SARS-CoV-2 for 30 min and then added to cell suspension. Cells in the plates were cultured for 2–3 days, and then exposed with MTT (3-[4,5-dimethyl-2-thiazolyl]-2,5-diphenyl-2H-tetrazolium bromide) (Nacalai Tesque). Cell viability was determined by measurement of absorbance at 560 nm and 690 nm. The concentration achieving 50% inhibition of cytopathic effect (effective concentration; EC_50_) was defined in GraphPad Prism version 8.4.3 (GraphPad Software) with a variable slope (four parameters). Non-treated cells were used as a control for 100% inhibition.

### Pharmacokinetics studies

Five-week old male Syrian hamsters (Japan SLC) were orally administered at 10, 30, and 100 mg/kg of S-217622 under the non-fasting conditions (n = 3, each). Dosing vehicles were 0.5% (w/v) methyl cellulose 400. Multiple blasma samples (0.1 mL) were collected over time from hamsters and plasma samples were stored in a freezer until analysis. Plasma concentrations of S-217622 were determined by liquid chromatography-tandem mass spectrometry (LC/MS/MS) system following protein precipitation with acetonitrile (MeCN). LC/MS/MS system equipped with a positive electrospray ionization (ESI) surface consisted of a API5000 (AB SCIEX, Framingham, MA, U.S.A.) and Nexera (Shimadzu Corporation, Kyoto, Japan). The multiple reaction monitoring (MRM) mode was selected and monitored the precursor ion (m/z 532.347) and product ion (m/z 145.042). The cone voltage and collision energy were set 90V and 55V, respectively. The column [YMC-Triart C18 (3 µm, 2.1 mm I.D.×50 mm), YMC Co., Ltd] was used and column temperature was maintained at 40°C for chromatographic separation of analytes. The mobile phases were 0.1% (v/v) formic acid in distilled water (mobile phase A) and MeCN (mobile phase B). The flow rate was 0.75 mL/min. The gradient condition was 30-65-95-95-30 (% of mobile phase B concentration) / 0 0.9-0.91-1.1-1.11-1.5 (min). Pharmacokinetic parameters of S 217622 in plasma were calculated by WinNonlin (Ver. 8.3, Certara, L.P.) based on a non compartment analysis with uniform weighting.

### Virus infection and treatment of hamsters

Five-week old male Syrian hamsters (Japan SLC) were intranasally inoculated with 5,000 pfu of SARS-CoV-2 in 200 μl of PBS under anesthesia with isoflurane inhalation. S-217622 was suspended in 0.5 % (w/v) methyl cellulose 400 and MPV was suspended in 10% polyethylene glycol 400 with 2.5% Cremophor RH40. For treatment, hamsters were orally administered twice daily with antivirals under anesthesia with isoflurane inhalation. Vehicle control hamsters were administered with 0.5 % (w/v) methyl cellulose 400. Body weights of animals were monitored daily. Nasal turbinates and lungs were collected from a subset of hamsters at 4 dpi and homogenized in 1ml or 5 ml of PBS with TissueRuptor (Qiagen), respectively. A part of the homogenate was subjected to plaque assays for virus titration. Total RNA was extracted from the homogenate with Direct-zol RNA MiniPrep kit and analyzed by qRT-PCR as mentioned above. To determine the titers of neutralizing antibody in hamsters, serum samples were collected from hamsters at 18 dpi and heat-inactivated at 56°C for 30 min. Virus neutralization assays were performed following the protocol as previously described^38^. The neutralization titer was defined as the reciprocal of the highest serum dilution that completely inhibited the cytopathic effect in Vero-TMPRSS2 cells.

For virus transmission between animals, one hamster per cage was inoculated with 5,000 pfu of SARS-CoV-2 Delta variant (infected hamster) and co-housed with three naïve hamsters (contact hamsters) in the same cage. Only contact hamsters were orally administered twice daily with antivirals from the time of co-housing. After 6 days, lung tissues were samples from contact hamsters and analyzed by virus titration and qRT-PCR as described above.

### Histopathological examination

Lung tissues at 4 dpi were fixed in 3.7% formaldehyde in PBS and embedded in paraffin. To detect viral RNA in the paraffin sections, ISH was carried out using an RNA scope 2.5 HD Red Detection kit (Advanced Cell Diagnostics) with antisense probe targeting the nucleocapsid gene of SARS-CoV-2 (Advanced Cell Diagnostics) as previously described^14^. Histopathological severity score of pneumonia was determined based on the percentage of alveolar inflammation in a given area of a pulmonary section collected from each animal in each group using the following scoring system: 0, no pathological change; 1, affected area (≤10%); 2, affected area (<50%, >10%); 3, affected area (≥50%); an additional point was added when pulmonary edema and/or alveolar hemorrhage was observed^14^. The total score for the five lobes was calculated for each animal.

### Light sheet microscopy

Samples for whole imaging analysis were prepared following the method of iDISCO as previously described with minor modifications^39^. Lung tissues at 4 dpi were fixed in 3.7% formaldehyde in PBS. The lungs were dehydrated by serial incubations in PBS, 50% methanol in PBS, 80% methanol in PBS, 100% methanol for 90 min each. Tissue bleaching was performed in a 9:1 mixture of 100% methanol and 30% H_2_O_2_ solution for overnight at 4°C. The lungs were rehydrated by serial incubations in 100% methanol twice, 80% methanol in PBS, 50% methanol in PBS, PBS for 60 min each. Tissue blocking was performed in PBSBT (20% Block Ace, 0.5% Triton X-100, 0.009% NaN_3_ in PBS) for 24 h. Tissues were then stained with anti-SARS-CoV-2 spike (1:2000, GTX635792, GeneTex) in PBSBT with 0.1% saponin for 4 days, followed by washing five times for 12 min each in PBS with 0.5% Triton X-100. Tissues were further incubated for 4 days with secondary antibody staining solution; Alexa Fluor Plus 647-conjugated anti-rabbit IgG (A32795, Invitrogen) diluted in PBSBT with 0.1% saponin and filtrated though a 0.45 μm syringe filter. After washing five times for 12 min each in PBS with 0.5% Triton X-100, the immunostained tissues were dehydrated by serial incubations in 50% methanol in PBS, 80% methanol in PBS, 100% methanol twice for 12 h each. Samples were then treated with dichloromethane for 40 min for removal of lipids, and then dibenzyl ether overnight. All incubation processes were conducted on an orbital shaker or a rotator. Fluorescence images were acquired by UltraMicroscope Blaze (Miltenyi Biotec) according to the manufactures’ instruction. Image data was converted and processed into 3D reconstruction by Imaris software (Oxford Instruments).

### Statistical analysis

Statistical significance was determined by two-tailed Mann-Whitney test (Extended Figs . 4a-f), one-way analysis of variance (ANOVA) with Dunnet’s test (Figs. 1d, 1e, 1h, 1i and Extended Fig. 3b), one-way ANOVA with Tukey’s test (Fig. 2h-k and 3g-j), Kruskal-Wallis test with Dunn’s multiple comparisons test (Fig. 1g, 2d-g, 2l, 2n, 2o, 3c-f, 4b and Extended Data Fig. 5b). All statistical tests were carried out using GraphPad Prism version 8.4.3 software.

## Acknowledgments

We thank National Institute of Infectious Diseases, Japan for providing SARS-CoV-2. We also thank Yuki Tachibana for providing S-217622; Katsuki Yokoo, Jun Sato, Tetsuya Miyano, and Yuto Unoh for providing nirmatrelvir; members in Section of Safety & DMPK Evaluation and Section of Analytical Chemistry & Bioanalysis 1 group, Shionogi TechnoAdvance Research Co., Ltd. for supporting pharmacokinetics analysis; Katsumi Maenaka and Basis for Supporting Innovative Drug Discovery and Life Science Research (BINDS) for supporting light sheet microscopy; Yuko Sato and Seiya Ozono for technical assistance. This work was supported by the Japan Agency for Medical Research and Development (AMED) under Grant numbers JP20fk0108509, JP21wm0125008, JP21wm0225003, JP21fk0108104; Japan Science and Technology Agency (JST) Moonshot R&D under Grant numbers JPMJMS2025; and the World-leading Innovative and Smart Education (WISE) Program from the Ministry of Education, Culture, Sports, Science, and Technology (MEXT), Japan under Grant numbers 1801; Hokkaido University COVID-19 Research Support Program; and COVID-19 Drug and Vaccine Development Donation.

## Author Contributions

M.S., K.U., Y.M., A.S. performed cell culture experiments, M.S., K.T., M.K., Y.I., H.K., T.A., S.T., S.K., H.S. performed animal infection experiments, S.Y. performed pharmacokinetics experiments, M.S., S.I., N.N., T. Suzuki, Y.O. performed pathological experiments, M.S., H.S. designed and supervised the study, H.N., T. Sanaki, T.K., T. Shishido provided critical resources, Y.O., A.S., H.S. obtained funding, M.S., W.W.H., H.S. wrote the draft. All authors reviewed and contributed to prepare the final version of manuscript.

## Data availability

All data are available within the article and extended data.

---

**Extended Data Figure 1.**
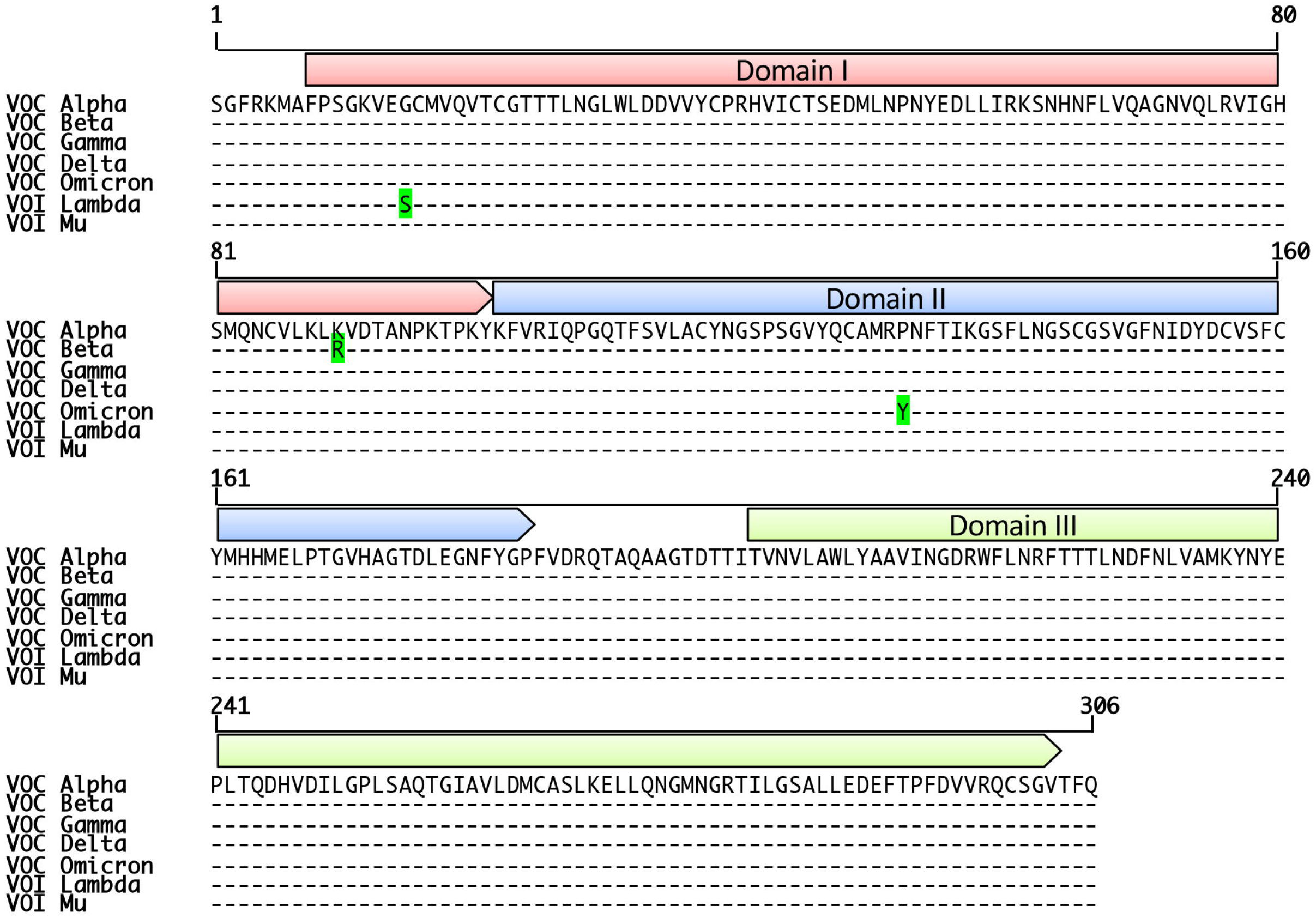
Multiple amino acid sequence alignments of M^pro^ from SARS-CoV-2 VOCs and VOIs. Multiple sequence alignment based on the entire length of M^pro^ from VOC Alpha (lineage B.1.1.7, GISAID: EPI_ISL_768526), Beta (lineage B.1.351, GISAID: EPI_ISL_1123289), Gamma (lineage P.1, GISAID: EPI_ISL_877769), Delta (lineage B.1.617.2, GISAID: EPI_ISL_2158617), Omicron (lineage BA.1, GISAID: EPI_ISL_7418017), VOI Lambda (lineage C.37, GISAID: EPI_ISL_4204973), and Mu (lineage B.1.612, GISAID: EPI_ISL_4470503). Dash (-) represents amino acids identical to M^pro^ of Alpha. Amino acids different from the consensus are colored in green.

**Extended Data Figure 2.**
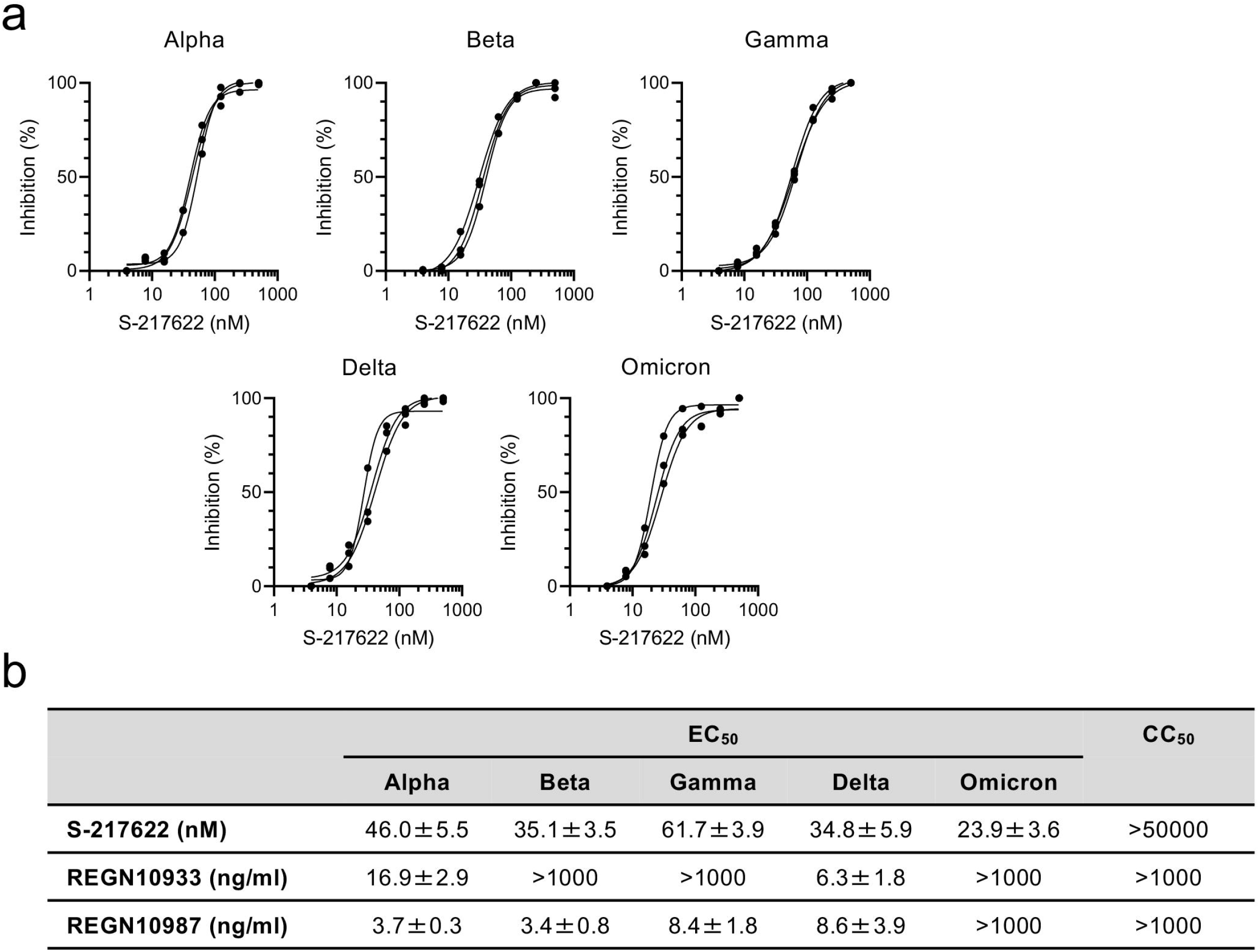
Antiviral potency of S-217622 against SARS-CoV-2 VOCs. Cytopathic effect (CPE) in 293T-hACE2-TMPRSS2 cells induced by SARS-CoV-2 infection was measured at 3 dpi by MTT assay. **a**, The inhibitory effect of S-217622 on SARS-CoV-2 VOCs was examined. Uninfected cells were used as a control for 100% inhibition. **b**, The 50% effective concentration (EC_50_) and cytotoxic concentration (CC_50_) of S-217622 and anti-SARS-CoV-2 neutralizing monoclonal antibodies (REGN10933 and REGN10987).

**Extended Data Figure 3.**
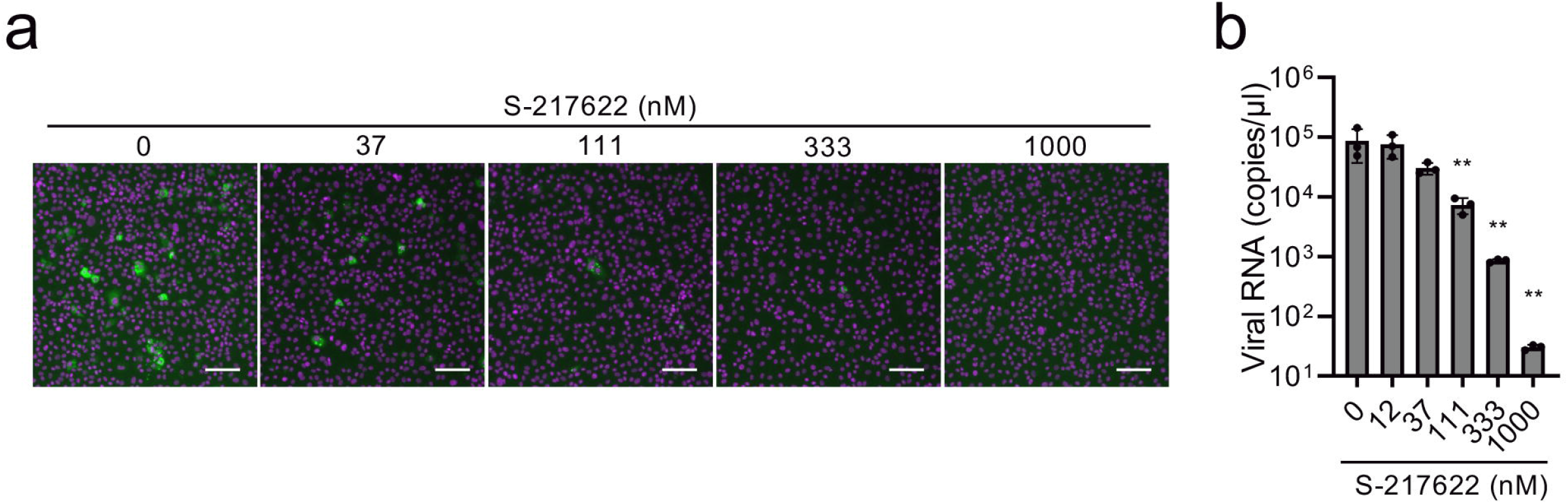
Susceptibility of SARS-CoV-2 Omicron variant to treatment with S-217622. **a**, Immunofluorescence staining of Vero-TMPRSS2 cells infected with the SARS-CoV-2 Omicron variant. Cells were inoculated with the SARS-CoV-2 Omicron variant and then cultured in the presence of S-217622 for 24 h. Cells were stained with anti-SARS-CoV-2 nucleocapsid antibody (green) and Hoechst 33342 (magenta). Scale bars, 100 μm. **b**, Viral RNA levels in the culture supernatant of Vero-TMPRSS2 at 24 hpi. The values shown are mean ± SD of triplicate samples. **p<0.01 by one-way ANOVA with Dunnet’s test.

**Extended Data Figure 4.**
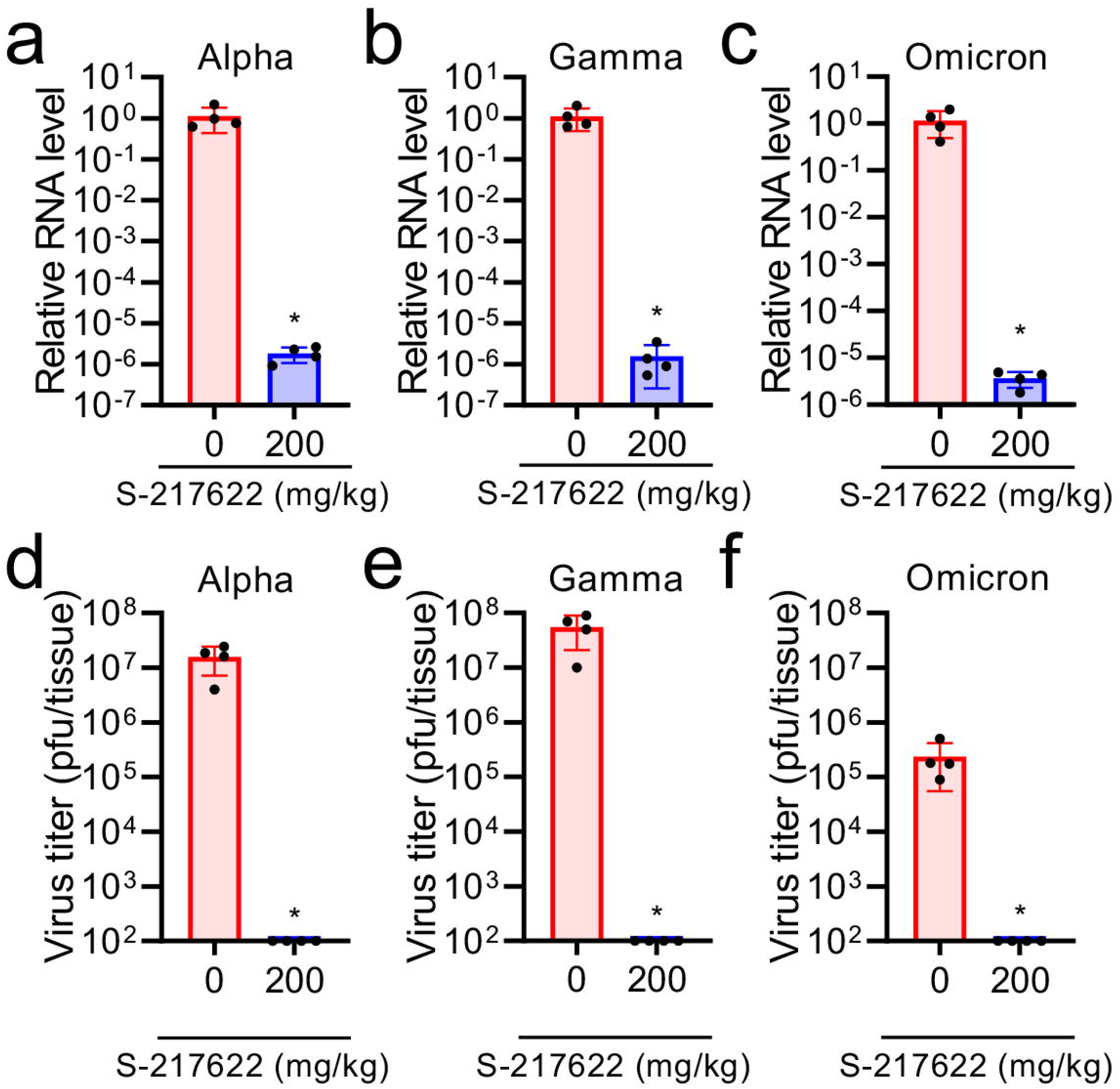
*In vivo* antiviral efficacy of S-217622 against SARS-CoV-2 VOCs. **a**-**f**, Hamsters were intranasally inoculated with 5,000 pfu of SARS-CoV-2 Alpha (**a**, **d**), Gamma (**b**, **e**), and Omicron (**c**, **f**) variants. The hamsters were treated with oral administration of S-217622 (200 mg/kg) or vehicle (0 mg/kg) twice a day from the time of inoculation to 3 dpi. Lung tissues were harvested at 4 dpi. **a**–**c**, Relative viral RNA levels in the lungs as compared with lungs from vehicle-treated hamsters were examined. Data were normalized to β-actin. **d**–**f**, Virus titers in the lungs were determined by plaque assay. The values shown are mean ± SD. (n = 4; *p<0.05 by two-tailed Mann-Whitney test.)

**Extended Data Figure 5.**
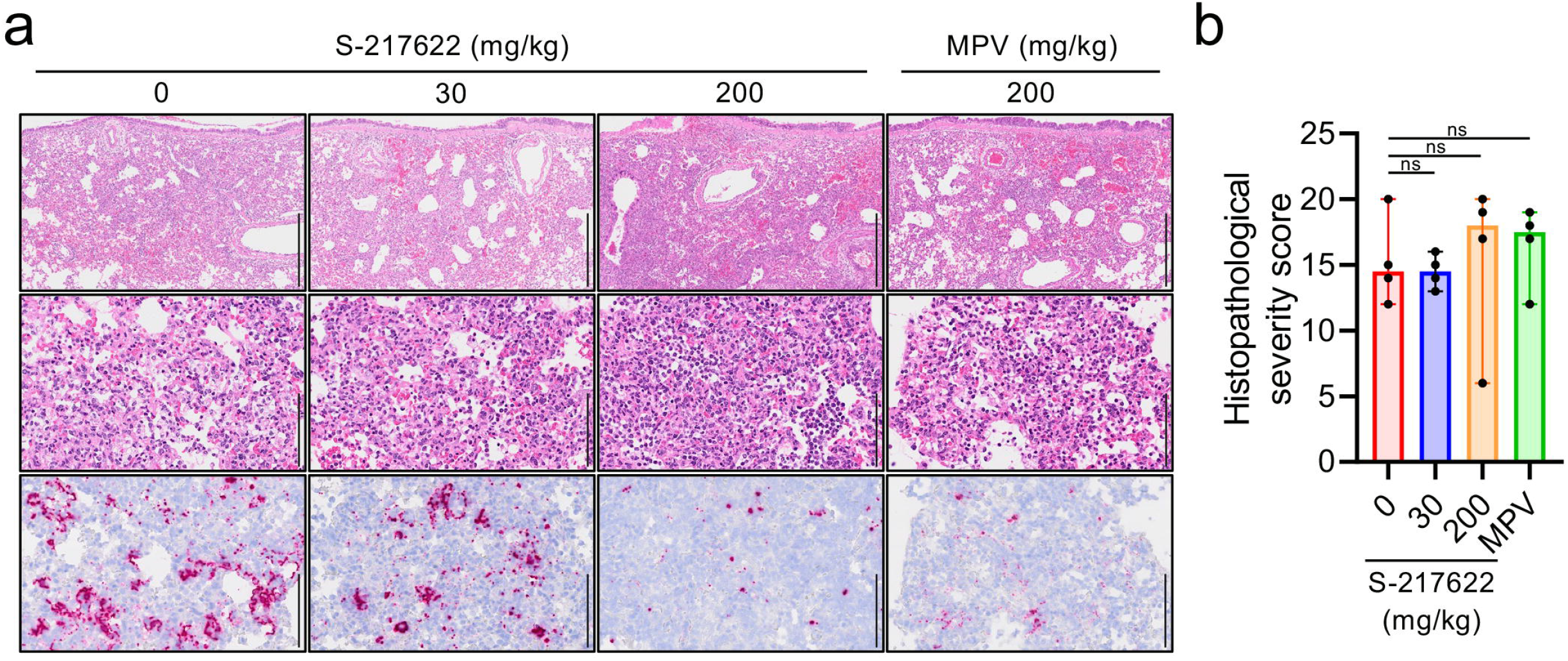
Histopathological findings in the lungs of SARS-CoV-2-infected hamsters that were administered drugs with therapeutic protocol. Hamsters were infected with the SARS-CoV-2 Delta variant and sacrificed at 4 dpi for histopathological examination. S-217622 (30 mg/kg or 200 mg/kg), MPV (200 mg/kg), or vehicle control (0 mg/ml) was administered from 1 to 3 dpi twice a day following the schedule shown in 3a. **a**, Representative histopathological images of lung sections obtained from the animals given antivirals are indicated above each panel (n = 4). Upper and middle panels, H&E staining. Lower panels, ISH targeting the nucleocapsid gene of SARS-CoV-2. Scale bars in middle and lower panels, 100 μm. **b**, Histopathological severity score of pneumonia based on the percentage of alveolitis in a given section. Data are shown as the median score ± 95% confidential interval with each dot representing the score of each animal. (n = 4; ns = not significant by Kruskal-Wallis test with Dunn’s test.)

**Extended Data Table 1.** Comparison of the potency of SARS-CoV-2 antivirals. Antiviral activity of the listed compounds against SARS-CoV-2 Delta was examined by CPE-based cell viability assay using (1) 293T-hACE2-TMPRSS2 cells, (2) Vero-TMPRSS2 cells, or (3) Vero-TMPRSS2 cells treated with a P-gp inhibitor (CP-100356).

|  | EC<sub>50</sub> against SARS-CoV-2 Delta |  |  |
| --- | --- | --- | --- |
|  | (1) 293T-hACE2-TMPRSS2 | (2) Vero-TMPRSS2 | (3) Vero-TMPRSS2 with CP-100356 |
| S-217622 (nM) | 26 ± 6.65 | 407 ± 21.3 | 69 ± 11.2 |
| Nirmatrelvir (nM) | 37 ± 5.30 | >1000 | 132 ± 32.0 |
| NHC (nM) | 538 ± 142 | 360 ± 49.4 | 677 ± 8.89 |
| Molnupiravir (nM) | 3675 ± 1093 | 3169 ± 217 | 2280 ± 118 |
| Remdesivir (nM) | <3.9 | >500 | 32 ± 4.52 |

**Extended Data Table 2.** Pharmacokinetic parameters of S-217622 in plasma after single oral administration of S-217622 in hamsters.

Extended Table 2
|  | S-217622 in plasma |  |  |
| --- | --- | --- | --- |
|  | 10 mg/kg | 30 mg/kg | 100 mg/kg |
| C<sub>max</sub> (μM) | 12.7 ± 2.2 | 61.6 ± 18.7 | 254 ± 24 |
| T<sub>max</sub> (h) | 0.5 ± 0.0 | 1.17 ± 0.76 | 2.67 ± 1.15 |
| t<sub>1/2,z</sub> (h)<sup>a</sup> | 4.46 ± 1.03 | 3.96 ± 0.74 | 3.43 ± 0.13 |
| AUC<sub>0-28h</sub> (μM·h) | 78.3 ± 11.0 | 415 ± 52 | 2480 ± 350 |
| AUC<sub>0-inf</sub> (μM·h) | 79.3 ± 10.5 | 417 ± 51 | 2500 ± 350 |

a: t<sub>1/2</sub>, 8-28hr

## Supplementary Information

**Supplementary Video 1.**

Related to Figure 4c. Reconstructed 3D image of viral antigen distribution in the lung of hamster at 4 dpi. The hamster was inoculated with SARS-CoV-2 Delta variant and treated with 0.5% methyl cellulose solution (vehicle for S-217622) from 1 dpi to 3 dpi.

**Supplementary Video 2.**

Related to Figure 4d. Reconstructed 3D image of viral antigen distribution in the lung of hamster at 4 dpi. The hamster was inoculated with SARS-CoV-2 Delta variant and treated with S-217622 (200 mg/kg) from 1 dpi to 3 dpi.

**Supplementary Video 3.**

Related to Figure 4e. Reconstructed 3D image of viral antigen distribution in the lung of hamster at 4 dpi. The hamster was inoculated with SARS-CoV-2 Delta variant and treated with MPV (200 mg/kg) from 1 dpi to 3 dpi.

## References

1. Tao, K. et al. SARS-CoV-2 Antiviral Therapy. Clin Microbiol Rev 34, e0010921, doi:10.1128/CMR.00109-21 (2021).

2. Fan, H., Lou, F., Fan, J., Li, M. & Tong, Y. The emergence of powerful oral anti-COVID-19 drugs in the post-vaccine era. Lancet Microbe 3, e91, doi:10.1016/S2666-5247(21)00278-0 (2022).

3. Owen, D. R. et al. An oral SARS-CoV-2 M^pro^ inhibitor clinical candidate for the treatment of COVID-19. Science 374, 1586–1593, doi:10.1126/science.abl4784 (2021).

4. Wahl, A. et al. SARS-CoV-2 infection is effectively treated and prevented by EIDD-2801. Nature 591, 451–457, doi:10.1038/s41586-021-03312-w (2021).

5. Rosenke, K. et al. Orally delivered MK-4482 inhibits SARS-CoV-2 replication in the Syrian hamster model. Nat Commun 12, 2295, doi:10.1038/s41467-021-22580-8 (2021).

6. Yang, H. & Rao, Z. Structural biology of SARS-CoV-2 and implications for therapeutic development. Nat Rev Microbiol 19, 685–700, doi:10.1038/s41579-021-00630-8 (2021).

7. Jin, Z. et al. Structure of M(pro) from SARS-CoV-2 and discovery of its inhibitors. Nature 582, 289–293, doi:10.1038/s41586-020-2223-y (2020).

8. Sourimant, J., Aggarwal, M. & Plemper, R. K. Progress and pitfalls of a year of drug repurposing screens against COVID-19. Curr Opin Virol 49, 183–193, doi:10.1016/j.coviro.2021.06.004 (2021).

9. Unoh, Y. et al. Discovery of S-217622, a Non-Covalent Oral SARS-CoV-2 3CL Protease Inhibitor Clinical Candidate for Treating COVID-19. bioRxiv, 2022.2001.2026.477782, doi:10.1101/2022.01.26.477782 (2022).

10. Shionogi & Co., L. Shionogi Presents Clinical Trial Results of the COVID-19 Therapeutic Drug S-217622. doi:https://www.shionogi.com/global/en/investors/ir-library/presentation-materials.html (2022).

11. Pizzorno, A. et al. Characterization and Treatment of SARS-CoV-2 in Nasal and Bronchial Human Airway Epithelia. Cell Rep Med 1, 100059, doi:10.1016/j.xcrm.2020.100059 (2020).

12. Boras, B. et al. Preclinical characterization of an intravenous coronavirus 3CL protease inhibitor for the potential treatment of COVID19. Nat Commun 12, 6055, doi:10.1038/s41467-021-26239-2 (2021).

13. Saito, A. et al. Enhanced fusogenicity and pathogenicity of SARS-CoV-2 Delta P681R mutation. Nature, doi:10.1038/s41586-021-04266-9 (2021).

14. Halfmann, P. J. et al. SARS-CoV-2 Omicron virus causes attenuated disease in mice and hamsters. Nature, doi:10.1038/s41586-022-04441-6 (2022).

15. Cao, Y. et al. Omicron escapes the majority of existing SARS-CoV-2 neutralizing antibodies. Nature, doi:10.1038/s41586-021-04385-3 (2021).

16. Harvey, W. T. et al. SARS-CoV-2 variants, spike mutations and immune escape. Nat Rev Microbiol 19, 409–424, doi:10.1038/s41579-021-00573-0 (2021).

17. Lou, F. et al. Understanding the Secret of SARS-CoV-2 Variants of Concern/Interest and Immune Escape. Front Immunol 12, 744242, doi:10.3389/fimmu.2021.744242 (2021).

18. Kilianski, A., Mielech, A. M., Deng, X. & Baker, S. C. Assessing activity and inhibition of Middle East respiratory syndrome coronavirus papain-like and 3C-like proteases using luciferase-based biosensors. J Virol 87, 11955–11962, doi:10.1128/JVI.02105-13 (2013).

19. Imai, M. et al. Syrian hamsters as a small animal model for SARS-CoV-2 infection and countermeasure development. Proc Natl Acad Sci U S A 117, 16587–16595, doi:10.1073/pnas.2009799117 (2020).

20. Sia, S. F. et al. Pathogenesis and transmission of SARS-CoV-2 in golden hamsters. Nature 583, 834–838, doi:10.1038/s41586-020-2342-5 (2020).

21. Kellam, P. & Barclay, W. The dynamics of humoral immune responses following SARS-CoV-2 infection and the potential for reinfection. J Gen Virol 101, 791–797, doi:10.1099/jgv.0.001439 (2020).

22. Yuan, L. et al. Gender associates with both susceptibility to infection and pathogenesis of SARS-CoV-2 in Syrian hamster. Signal Transduct Target Ther 6, 136, doi:10.1038/s41392-021-00552-0 (2021).

23. Zhang, L. et al. -Ketoamides as Broad-Spectrum Inhibitors of Coronavirus and α Enterovirus Replication: Structure-Based Design, Synthesis, and Activity Assessment. J Med Chem 63, 4562–4578, doi:10.1021/acs.jmedchem.9b01828 (2020).

24. Cox, R. M. et al. Oral prodrug of remdesivir parent GS-441524 is efficacious against SARS-CoV-2 in ferrets. Nat Commun 12, 6415, doi:10.1038/s41467-021-26760-4 (2021).

25. Yuan, S. et al. Clofazimine broadly inhibits coronaviruses including SARS-CoV-2. Nature 593, 418–423, doi:10.1038/s41586-021-03431-4 (2021).

26. Jayk Bernal, A. et al. Molnupiravir for Oral Treatment of Covid-19 in Nonhospitalized Patients. N Engl J Med, doi:10.1056/NEJMoa2116044 (2021).

27. Max, K. Merck’s COVID pill loses its lustre: What that means for the pandemic. Nature (2021).

28. Zhou, S. et al. -d-N4-hydroxycytidine Inhibits SARS-CoV-2 Through Lethal β Mutagenesis But Is Also Mutagenic To Mammalian Cells. J Infect Dis 224, 415–419, doi:10.1093/infdis/jiab247 (2021).

29. Administration, U. F. a. D. (2021).

30. Heskin, J. et al. Caution required with use of ritonavir-boosted PF-07321332 in COVID-19 management. Lancet 399, 21–22, doi:10.1016/S0140-6736(21)02657-X (2022).

31. Thorne, L. G. et al. SARS-CoV-2 sensing by RIG-I and MDA5 links epithelial infection to macrophage inflammation. EMBO J 40, e107826, doi:10.15252/embj.2021107826 (2021).

32. Tay, M. Z., Poh, C. M., Rénia, L., MacAry, P. A. & Ng, L. F. P. The trinity of COVID-19: immunity, inflammation and intervention. Nat Rev Immunol 20, 363–374, doi:10.1038/s41577-020-0311-8 (2020).

33. Feuillet, V., Canard, B. & Trautmann, A. Combining Antivirals and Immunomodulators to Fight COVID-19. Trends Immunol 42, 31–44, doi:10.1016/j.it.2020.11.003 (2021).

34. Sasaki, M. et al. SARS-CoV-2 variants with mutations at the S1/S2 cleavage site are generated in vitro during propagation in TMPRSS2-deficient cells. PLoS Pathog 17, e1009233, doi:10.1371/journal.ppat.1009233 (2021).

35. Sasaki, M. et al. Air-liquid interphase culture confers SARS-CoV-2 susceptibility to A549 alveolar epithelial cells. Biochem Biophys Res Commun 577, 146–151, doi:10.1016/j.bbrc.2021.09.015 (2021).

36. Shirato, K. et al. Development of Genetic Diagnostic Methods for Detection for Novel Coronavirus 2019(nCoV-2019) in Japan. Jpn J Infect Dis 73, 304–307, doi:10.7883/yoken.JJID.2020.061 (2020).

37. Zivcec, M., Safronetz, D., Haddock, E., Feldmann, H. & Ebihara, H. Validation of assays to monitor immune responses in the Syrian golden hamster (Mesocricetus auratus). J Immunol Methods 368, 24–35, doi:10.1016/j.jim.2011.02.004 (2011).

38. Sasaki, M. et al. SARS-CoV-2 Bearing a Mutation at the S1/S2 Cleavage Site Exhibits Attenuated Virulence and Confers Protective Immunity. mBio 12, e0141521, doi:10.1128/mBio.01415-21 (2021).

39. Tomris, I. et al. 3D visualization of SARS-CoV-2 infection and receptor distribution in Syrian hamster lung lobes display distinct spatial arrangements. bioRxiv, 2021.2003.2024.435771, doi:10.1101/2021.03.24.435771 (2021).

